# Yo-yo dieting deregulates feeding behavior in mice via the induction of durable gut dysbiosis

**DOI:** 10.1101/2024.12.02.625705

**Authors:** Mélanie Fouesnard, Adélie Salin, Sandy Ribes, Magali Monnoye, Gaëlle Champeil-Potokar, Marie-Sabelle Hjeij, Gwénaëlle Randuineau, Léa Le Gleau, Selma Ben Fradj, Catherine Philippe, Alexandre Benani, Isabelle Denis, Véronique Douard, Gaëlle Boudry

**Affiliations:** Institut Numecan, INRAE, INSERM, Univ Rennes, Rennes, France; Institut MICALIS, INRAE, AgroParisTech, Université Paris-Saclay, Jouy-en-Josas, France; UMR PNCA, INRAE, AgroParisTech, Université Paris-Saclay, Palaiseau, France; Centre des Sciences du Goût et de l’Alimentation, UMR CNRS U6265 INRAE U13241 Université de Bourgogne, Dijon, France

**Keywords:** Microbiota-gut-brain axis, eating behavior, hedonic appetite

## Abstract

**Background & Aims:** Alternating periods of excessive and restrained eating results in weight cycling, a known risk factor for eating behavior dysregulation such as binge eating. Diet alternation also induces changes in intestinal microbiota composition. We tested the hypothesis that recurrent diet alternation alters hedonic feeding regulation by changing either or both intestinal microbiota and brain homeostasis in mouse.

**Methods:** C57BL/6 mice underwent 3 cycles of 1 week of western diet (WD, 45% kcal from fat) separated by 2 weeks of chow diet (CYCL group) or staid under chow diet (CTRL group). Food intake was monitored after each dietary change. Striatum, hypothalamus, brainstem and caecal content were collected before the third WD introduction in CYCL mice and in CTRL mice. Microbiota transfer from CYCL or CTRL mice into naive recipient mice was performed to investigate whether gut microbiota per se could explain differences in eating behavior during weight cycling.

**Results:** Diet alternation in CYCL mice resulted in weight cycling, with enhanced weight gain upon each WD feeding phase. CYCL mice increased their energy intake specifically during the first hours following WD re-introduction, reminiscent of binge-eating episodes. Expression of reward-related genes in the striatum and thickness of the astro-glial barrier in the brain stem were enhanced in CYCL compared to CTRL mice. Diet alternation also induced caecal dysbiosis in CYCL mice. Gut microbiota transfer from CYCL mice to naive recipient mice recapitulated the altered eating behavior upon WD exposure.

**Conclusions:** Alternation between high-energy and standard diet durably remodels the gut microbiota and the brain towards a profile associated with an increase in hedonic appetite. Using gut microbiota transfer, we established that this microbiota signature affects hedonic feeding regulation. These results open the ways to microbiota-targeted strategies to prevent development of eating disorders in weight cycling patients.

## Introduction

The rise in overweight and obesity together with the social pressure for thinness increase the prevalence of dieting. Restrictive dieting practices have risen over the last few decades in Western countries, reaching 27% of the French population (1) and affecting up to 40% of men and 57% of women in the US during the 2003-2008 period (2). However, long-term studies report that restrictive dieting is ineffective in maintaining long-term weight reduction. Consequently, individuals following these restrictive dietary habits experience weight cycling characterized by repeated episodes of weight loss and subsequent excessive weight regain (3). In the US, this ‘yoyo’ effect affects between 20% and 35% of men and between 20% and 55% of women (4). Weight cycling probably reflects physiological adaptations that would originally contributed to ensure increased survival capacity during repeated periods of feast and famine. However in our modern societies it could lead to adverse health consequences (5). In terms of metabolic consequences, restrictive dieting is associated with changes in energy partitioning between fat and lean mass, alterations in cellular energetic efficiency and modifications of insulin and leptin sensitivity (5). Overall, a negative energy balance leads to an adaptative decrease of total daily energy expenditure (6) that is accompanied by hormonal changes (7) that both can last for years in case of profound weight loss (8,9). These adaptations turn unfavorable when energy restriction is discontinued, impeding efforts to maintain body weight and favoring weight regain. Besides hormonal and metabolic disturbances, strong weight loss is accompanied by an increased orexigenic drive (10). Several clinical studies suggest that repeated dieting and weight cycling predispose to disordered eating, including binge eating. In a multisite US study enrolling 1875 obese patients, 51% of binge-eating disorders (BED)-positive patients had gained or lost weight five times or more in their lifetime, compared with 27% in BED-negative patients (11). Similarly, cross-sectional studies have shown a positive relationship between weight cycling and binge eating: the greater the number of weight loss attempts, the greater the occurrence of binge eating (12,13). However, these studies were cross-sectional or retrospective, preventing any conclusion regarding the causal relationship between dieting and dysregulation of eating behavior. Therefore, preclinical models of binge eating and weight cycling are instrumental for studying such associations. While short-time access (1 or 2 hours per day) to a hypercaloric palatable diet induces an escalation of palatable diet consumption in rodents, reminiscent of binge eating, this model does not recapitulate weight cycling. In contrast, the alternation of extended periods (days to weeks) of a hypercaloric and a standard diet is associated with weight cycling as well as metabolic and behavioral adaptations (14,15). Interestingly, these studies have also underscored the key role of gut microbiota in post-dieting weight regain in obesity, attributing this effect to changes in energy expenditure.

Since the gut microbiome plays a crucial role in energy metabolism (by the conversion of dietary fiber into short-chain fatty acids (SCFAs)), and modulation of inflammation (16).Since it profoundly impacts many physiological systems and behaviors (17), including eating behaviors (18–21) we sought to investigate whether weight cycling was associated with deregulation of food intake driven by a peculiar gut microbiota signature. Noteworthy, we investigated the impact of weight cycling before the installation of obesity to avoid the confounding effects of significant obesity that is associated with profound modifications of energy expenditure and metabolic alterations such as hyperglycemia, hyper-leptinemia, or insulin resistance, all affecting food intake regulation. Hence, we exposed mice to three successive dietary switches between a standard rodent diet and a western diet (WD). Food intake was monitored for the duration of the procedure. To assess the consequences of yoyo dieting on the microbiota-brain axis, brain and microbiota analysis were performed after the three dietary switches. To causally link gut microbiota signatures to changes in eating behavior, we performed microbiota transfer into naive mice and determined the impact on eating behavior.

## Methods

### Animals

The present protocols received written agreement from the local ethic committees (file no. APAFIS#40563-2023013010327051v2 for experiments 1 and 2, and APAFIS#2020051821292405 for experiment 3). The experiments comply with the ARRIVE guidelines and have been carried out in accordance with the EU Directive 2010/63/EU for animal experiments. For all experiments, 7-week old mice were housed individually at arrival and fed a regular chow diet (chow, ref V1124-000, SSniff, Germany) *ad libitum* under a 12:12h light/dark cycle (22 ± 2°C). They were acclimatized to the animal facility for 1 week. Western Diet (WD, ref RD12451, Research diet, USA) was introduced at the beginning of the light cycle (resting period) for each cycle and mice were euthanized during the first half of the light cycle.

Experiment 1. C57Bl6/N male mice (Janvier labs, France) of similar body weights were divided into two groups: CYCL mice underwent 3 cycles of 1 week of WD feeding each followed by 2 weeks of chow diet feeding (n = 6). All diets were provided *ad libitum*. CTRL mice received chow diet during the whole experiment (n = 6). Food intake was measured during the first 8 hours following WD introduction (from 9am to 5pm) and the following 16 hours (5pm to day+1 9am) for each of the 3 WD cycles. Food intake and body weight were also measured daily during all the experiment. Fecal samples were collected at different time points along the experiment. Experimental protocol is summarized in **Fig. 1A**. Food consumption was expressed as caloric intake normalized to body weight.

**Figure 1.**
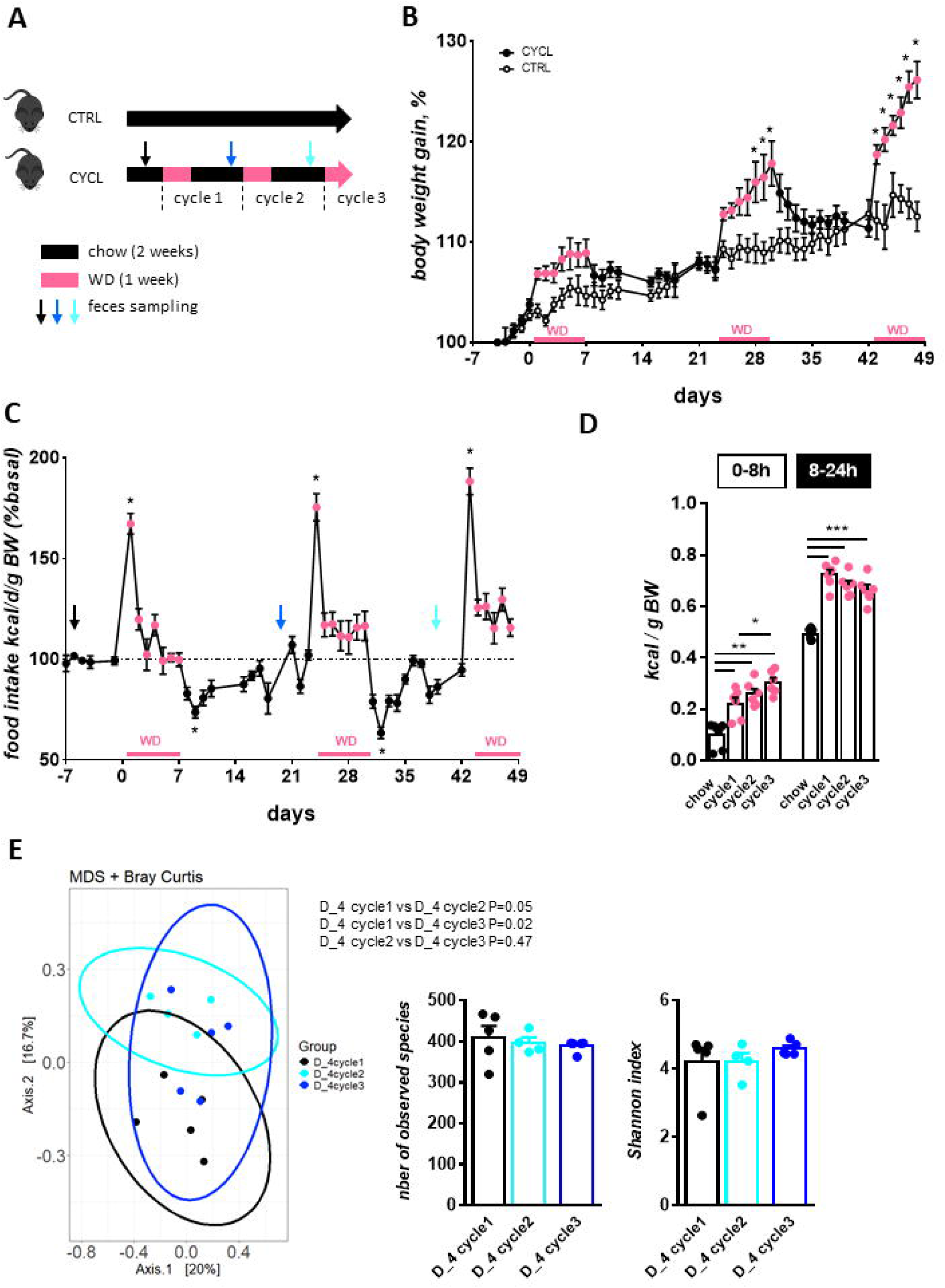
CYCL mice have altered regulation of energy intake upon WD introduction and display fecal dysbiosis. **(A)** Experimental design: Mice were submitted to cycles of one week of western diet (WD) followed by 2 weeks of chow diet (CYCL mice) or left under chow diet (CTRL). WD was introduced 2 hours after the onset of the light period. **(B)** Change in body weight of CTRL and CYCL mice during the whole protocol. **(C)** Daily caloric intake of CYCL mice during the whole protocol. **(D)** Caloric intake during the first 8 hours and following 16 hours of CYCL mice upon WD introduction at each cycle or under chow (average chow intake of each day before WD introduction). **(E)** Mouse fecal microbiota (Bray Curtis distance and α-diversity indices) 4 days before WD introduction at each cycle. For figure 1A to 1D, n=6 mice / group, * P<0.05 (Mann-Whitney test). For figure 1E, n = 4-5 mice per group, PERMANOVA test was performed on the Bray–Curtis dissimilarity matrix and Kruskal-Wallis test was performed on alpha diversity index. Each dot represents one mouse. Pink solid dots depict data when CYCL mice were under WD. Bars are means ± SEM.

Experiment 2 was conducted in two sets. In the first set, C57Bl6/N male mice (Janvier labs, France) were divided into 2 groups of similar body weights: CTRL mice were fed the chow diet *ad libitum*. CYCL mice underwent 2 cycles of 1 week of WD feeding each followed by 2 weeks of chow diet and were euthanized at the end of the 2^nd^ chow period (n = 7-8 for each group). Experimental protocol is summarized in **Fig. 2A**. Food intake was measured daily during the whole experiment. Caecal content and striatum were stored at −80°C after sampling for later 16S RNA and gene expression analysis respectively. Brainstems were dissected, fixed in 10% neutral buffered formalin (formaldehyde 4%, Diapath) for 24h at 4°C, cryoprotected with sucrose (30% in PBS), frozen in isopentane (−40◦C), then coronal sections were performed on a cryostat (Leica CM3050S) and collected on slides for immunohistochemistry. The second set followed the same design with n = 7 for each group). The hypothalamus and area postrema (AP) of the brainstem were dissected and stored at −80°C after sampling for later gene expression analysis.

**Figure 2.**
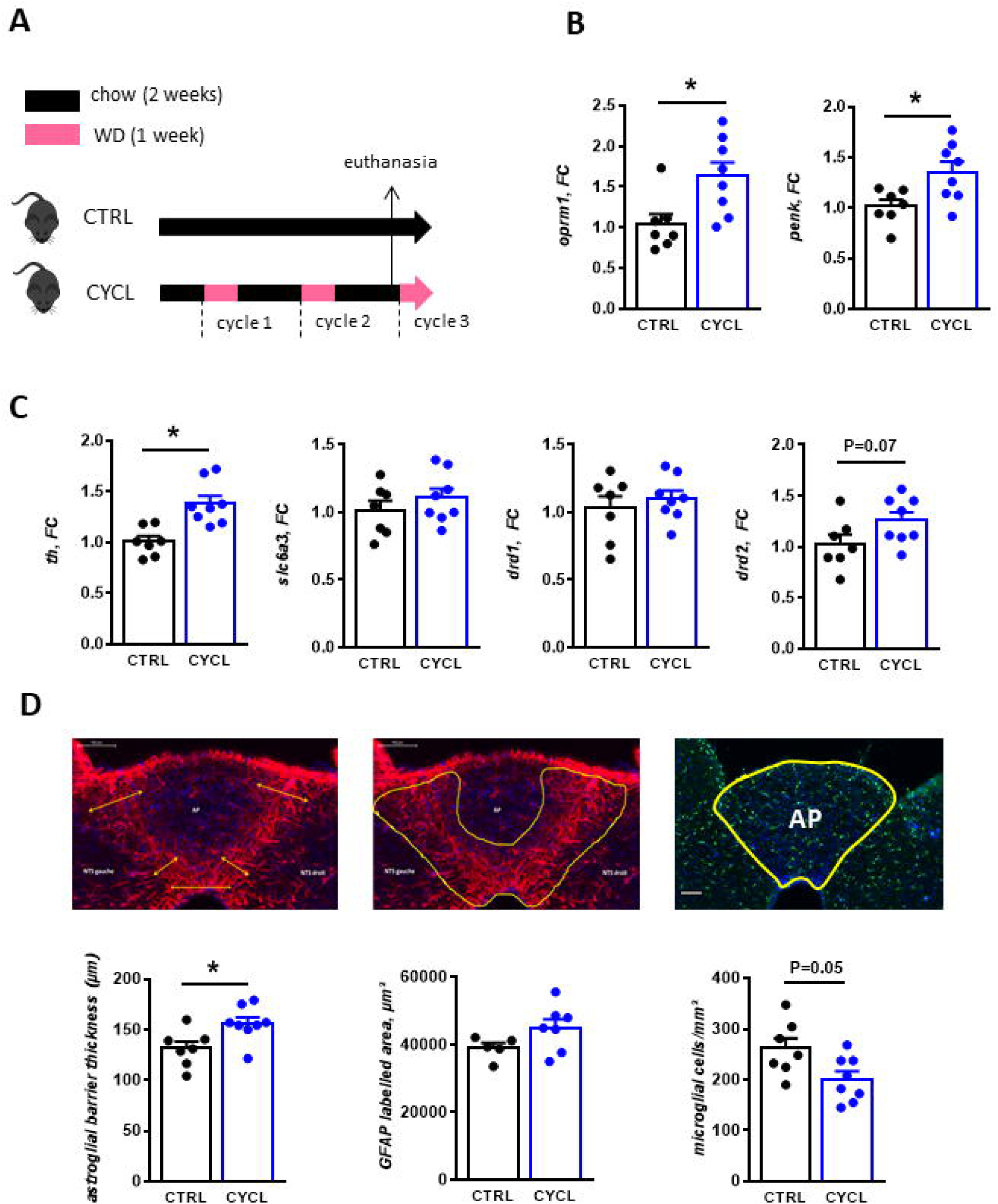
CYCL mice under chow diet display heightened reward phenotype in the striatum and altered barrier properties in the dorso-vagal complex. **(A)** Experimental design: Mice were submitted to cycles of one week of western diet (WD) followed by 2 weeks of chow diet (CYCL mice) or were left under chow diet (CTRL). They were euthanized while fed chow diet one day before the beginning of the third WD cycle. Age-matched CTRL mice fed only chow diet were euthanized at the same time. **(B)** Relative expression of genes involved in the opioïdergic system (*oprm1:* Opioid Receptor Mu 1 and *penk*: proenkephalin) in the striatum of CTRL and CYCL mice under chow diet. **(C)** Relative expression of genes involved in the dopaminergic system (*th:* tyrosine hydroxylase, *slc6a3*: dopamine transporter 1, *drd1:* dopamine receptor 1 and *drd2*: dopamine receptor 2) in the striatum of CTRL and CYCL mice under chow diet. **(D)** Astroglial barrier thickness between the area postrema (AP) and the nucleus tractus solitarius (NTS), astroglial cell (GFAP labelled) spreading between AP and NTS and microglial cell density in the AP of CTRL and CYCL mice under chow diet. Representative images are presented above graphs. n=7-8 mice / group, * P<0.05 (Mann-Whitney test). Each dot represents one mouse. Bars are means ± SEM.

Experiment 3. *Donor mice (n = 6 per group)*: C57Bl6/N male mice (Envigo, France) were divided into 2 groups of similar body weights: CTRL mice were fed a chow diet during all the experiment and CYCL mice underwent 2 cycles of 1 week of WD consumption each followed by 2 weeks of chow diet consumption. Mice were euthanized at the end of the 2^nd^ chow period. *Preparation of the inoculum*: Caecal contents were sampled from donor mice at the time of euthanasia, diluted at a 1:50 (mass:volume) ratio in maltodextrin-trehalose diluent (22), aliquoted and transferred to recipient mice or stored at −80°C. *Recipient mice (n = 12 per group)*: C57Bl6/N male mice (Envigo, France) were gavaged 5 times within a 2-hour period with 200µL of a polyethylene glycol (PEG) solution (Macrogol, 4000, 425g/L in sterile water) to deplete their gut microbiota. They were then divided into 2 groups of weight-matched microbiota-transferred (MT) mice (MT-CTRL and MT-CYCL). Four hours after the last PEG gavage, they were colonized by intra-gastric gavage of 200 μL of inoculum prepared from CTRL and CYCL mice, respectively. One donor mouse colonized 2 recipient mice. The next two days after transfer, recipient mice were gavaged with 200µL of frozen/thawed of the same inoculum. Body weight and food intake of recipient mice was monitored every week. Three weeks after inoculation, half of the recipient mice were euthanized to collect caecal contents. The daily food intake of the second half of the mice was monitored during the last 24h under chow diet and during 5 days after a switch to WD. Every day food intake was measured between 9am to 5pm (0-8h) and during the following 16 hours (8-24h). Experimental protocol is summarized in **Fig. 4A**.

### Determination of caecal microbiota composition

Total DNA was extracted from caecal content samples using the ZR fecal DNA Miniprep (Ozyme). The V3-V4 region of DNA coding for 16SrRNA was amplified using the following primer: CTTTCCCTACACGACGCTCTTCCGATCTACTCCTACGGGAGGCAGCAG (V3F) and GGAGTTCAGACGTGTGCTCTTCCGATCTTACCAGGGTATCTAATCC (V4R), Taq Phusion (New England Biolabs) and dNTP (New England Biolabs) during 25 cycles (10 s at 98°C, 30s at 45°C, 45s at 72°C). Purity of amplicons was checked on agarose gels before sequencing using Illumina Miseq technology, performed at the Genotoul Get-Plage facility (Toulouse, France). Sequences are available under https://doi.org/10.57745/OK3ZHQ. Raw sequences were analyzed using the bioinformatic pipeline FROGS (23). Amplicon Operational Taxonomic Unit (OTU) were clustered using Swarm with parameter d = 1 and chimeras were filtered following FROGS version 4.1 guidelines. Assignation was performed using 16S_REFseq_Bacteria_20230726 database. OTU with abundances lower than 0.0005% of the total read set were removed prior to analysis. Subsequent analysis were done using the phyloseq R packages (24). Samples were rarefied to even sampling depths before computing within-samples compositional diversities (observed richness and Shannon index) and between-samples compositional diversity to performed Multidimensional Scaling (MDS) analysis using Bray-Curtis or Jaccard dissimilarity. Raw, unrarefied OTU counts were used to produce relative abundance graphs.

### Determination of caecal SCFA levels

SCFA caecal levels were determined as previously described (10.1017/S0007114508978284). Briefly, SCFA were water-extracted and proteins were precipitated with phosphotungstic acid. Supernatant (containing SCFA) was analyzed in duplicate using gas-liquid chromatography (Auto-system XL; Perkin Elmer, Saint-Quentin-en-Yvelines, France). Samples were analysed in duplicate. Peaks were integrated using the Turbochrom v6 software (Perkin Elmer, Courtaboeuf, France).

### Brainstem glial immunohistochemistry

Immunohistochemistry (IHC) was performed on 20-μm thick serial horizontal brainstem sections containing the AP and tractus solitarius nucleus (NTS) to evaluate microglial and astroglial morphology. Non-specific sites were blocked by incubation with bovine serum albumin (BSA) (2% in PBS 0.1M; Triton X-100 0.3%) for 2 hours at room temperature. Astrocytes were labelled for GFAP (Glial fibrillary acidic protein) by incubating the slices for 1 hour with a monoclonal Cy3-anti-GFAP antibody (Sigma, France). Microglia were labelled for Iba1 (Ionized Calcium-binding Adapter Molecule 1) by incubating the slices for 48 hours at 4°C with a rabbit monoclonal anti-Iba1 antibody (Abcam, France), rinsed 3 times, then incubated for 24 hours at 4°C with a secondary Alexa 488-anti-rabbit antibody (Life Technologies, France). All antibodies were diluted 1:1000 in PBS-Triton-BSA 0.2%. Cell nuclei were labelled with bis-benzimide (0.5 μg/mL, Hoechst 33258, Sigma, France). Preparations were mounted in fluoroshield mounting medium (Sigma, France). IHC images from brain stem sections were acquired on a high capacity digital slide fluorescence scanner (Pannoramic SCAN P150, 3D HISTECH) at 20x resolution, using the extended focus function scanner to generate images with extended depth of field from acquired Z-stacks (11 images, 1 µm interval, 10 µm total thickness).

Iba1-labelled microglial cells within the AP were manually counted on 1 slice per mouse. In the dorsal vagal complex (DVC) sections, the deployment of the astroglial barrier was quantified by measuring (i) the total GFAP area delineating the border between the AP and the NTS on images converted in binary format using an automatic thresholding method (Threshold: “Li” algorithm) on Image J software as previously described (26), and (ii) 5 widths representative of the thickness of the barrier : dorsal-left, ventral left, dorsal right, ventral right, ventral central. Measurements were performed on 1 slice per mouse, each slice being identically positioned to allow comparisons between groups. All measurements were performed by the same experimenter who was blind to the groups.

### Striatum, hypothalamus and area postrema gene expression

Hypothalamus and AP total RNA were extracted using miRVANA RNA isolation kit (Ambion, Austin, TX, USA), following manufacturer’s instructions as described previously (25). Reverse transcription was performed from 1.8µg of total RNA from the hypothalamus and striatum and 1µg of total RNA from the AP using the cDNA high capacity reverse transcription kit (Applied Biosystems, Wattham, Massachusetts, USA) following manufacturer’s instructions. In the striatum, cDNAs diluted 1/7 were amplified by real time PCR using commercially available FAM-labeled Taqman® probes (ThermoFisher Scientific, France), [Opioid Receptor Mu 1 (*oprm1,* Mm01188089 ml), Proenkephalin (*penk*, Mm01212875_m1), Tyrosine Hydroxylase (*th*, Mm00447557_m1), dopamine receptor D1 (*drd1*, Mm02620146_s1), dopamine receptor D2 (*drd2,* Mm00438545_m1), Solute Carrier Family 6 Member 3 (*slc6a3*, Mm00438388_m1)]. Ribosomal protein lateral stalk subunit P0 (*rplp0,* Mm00725448_s1) and β-actin (*actb,* Mm02619580_g1) were used as reference genes.

In the hypothalamus and AP, gene expression of a total of 48 known candidate genes was studied and analyzed using real time TaqMan Low-Density Array (TLDA). Pre-designed 384-well format gene expression assays were used for the experiments (ThermoFisher Scientific, France). The primers and probe for each assay were preloaded onto the designated well. Forty-three candidate genes involved in regulation of food intake, brain plasticity, blood-brain barrier and microglia, astrocytes & inflammation were selected. In addition, five genes, *actb*, *rplp0*, glyceraldehyde-3-phosphate dehydrogenase (*gapdh*), phosphoglycerate kinase 1 (*pgk1*) and 18S ribosomal RNA (*18s*), were selected for spotting on the TLDA cards as endogenous controls. The genes and their functional relevance are listed in Suppl. Table 1.

For both, real time PCR and TLDA, the relative expression levels of target genes were calculated using the 2-ΔΔCT method considering the geometric mean of reference genes expression levels for determining the ΔCT for each sample and the CTRL as the reference group to determine the ΔΔCT.

### Statistics

Data between groups were compared using Kruskal-Wallis test (experiment 1) and Mann-Whitney test (experiments 2 & 3) on Prism GraphPad v.7.00 (GraphPad Software, San Diego, USA).

For microbiota analysis, alpha diversity index (Observed species and Shannon index) were analyzed using Kruskal-Wallis test (experiment 1) and Mann-Whitney test (experiments 2 & 3). A permutational multivariate analysis of variance (PERMANOVA) test was performed on the Bray–Curtis and Jaccard matrices using 9999 random permutations and at a significance level of 0.05. Phylum and family relative abundances were compared using a Mann-Whitney test.

## Results

### Weight cycling leads to transient loss of control over WD consumption

To examine the impact of weight cycling on eating behavior, CYCL mice underwent 3 cycles of 1 week of WD each followed by 2 weeks of chow diet feeding (**Fig. 1A**). In response to WD, CYCL mice gained significantly more body weight than CTRL mice during cycles 2 and 3 (**Fig. 1B**). CYCL mice lost weight each time they switched back to the chow diet, resulting in similar body weight than that of CTRL mice at the beginning of each new WD cycle (**Fig. 1B**). Moreover, the WD-induced body weight gain increased with the number of cycles (weekly change in body weight, cycle 1: 2.1 ±1.5%, cycle 2: 5.0 ±1.8%, cycle 3: 7.4 ±1.3%, P<0.05 for each comparison), reminiscent of the yoyo effect.

In response to each WD introduction, mice showed a 24-h hyperphagic phase followed by normalization of their energy intake within the five following days (**Fig. 1C**). When CYCL mice switched back to the chow diet, their daily energy intake decreased significantly when compared to the intake on the last day of WD (**Fig 1C**). The energy intake during the 24-h hyperphagic phase upon WD introduction did not differ among WD cycles. However, when we evaluated the WD consumption pattern during the first hours following WD introduction, we observed a major increased intake during the first 8 hours following WD reintroduction (**Fig. 1D**). Noteworthy, WD was introduced in home cages 2 hours after the onset of the light period, when mice are usually at rest, suggesting that CYCL mice displayed a period of loss of control over food intake during the light phase. Moreover, WD consumption during these first 8 hours increased with the number of cycles, with a 32% increase between cycles 1 and 3 (**Fig. 1D**). During the following 16 hours, there was a tendency (P=0.08) for a decrease in energy intake with increasing number of cycles (**Fig. 1D**) likely explaining the lack of difference in total 24-h calorie intake between WD cycles.

Yoyo dieting has already been shown to induce intestinal dysbiosis, even after mice have returned to chow diet (14). Therefore, we evaluated if the microbiota of CYCL mice was also altered by the alternance of WD and chow diets. Fecal samples were collected 4 days before each WD cycle when CYCL mice were fed chow diet and had returned to a body weight and energy intake similar to those of CTRL mice. As expected, recurrent exposure to WD induced fecal dysbiosis, as shown by the principal coordinate analysis with Bray Curtis distance (**Fig. 1E**). However, no difference was observed for the number of observed species nor the Shannon index between cycles (**Fig. 1E**).

### Weight cycling induces remodeling of central structures involved in food intake control

Eating behavior is regulated by the integration of signals originating from the gut (nutrients, bacterial metabolites) at different levels of the central nervous system, in particular in the DVC, the hypothalamus and the striatum. Therefore, we evaluated the impact of recurrent exposure to WD on the DVC, hypothalamus and striatum, by collecting samples from CYCL-chow mice the day before WD was reintroduced for the third time and comparing them to age-matched CTRL mice (**Fig. 2A**).

In the striatum, dietary alternation induced alterations in both the opioidergic and dopaminergic systems in CYCL mice compared to CTRL mice, both under the chow diet. These changes were characterized by a significant increase in gene expression of the opioid receptor mu 1 (*oprm1*) and proenkephalin (*penk*) (**Fig. 2B**) as well as tyrosine hydroxylase (*th*), the rate-limiting enzyme in the biosynthesis of catecholamines, including dopamine. Additionally, there was a trend towards increased expression of the dopamine receptor 2 (*drd2*) (**Fig. 2C**) in CYCL mice. Expression assay in the whole hypothalamus did not reveal any change in the relative expression of genes involved in regulation of food intake, plasticity, blood-brain barrier, astrocyte and microglia markers or inflammatory markers following dietary alternance (**Suppl. Table 1**).

In the DVC, the AP hosts a rich population of microglia and is bordered by a thick barrier of astroglial cells that may control the transfer of circulating molecules from the AP to the blood-brain barrier-protected nucleus of the DVC. These glial populations have been shown to react to WD (27,28). In our study, the deployment of the astroglial barrier between the AP and the NTS was increased, showing a larger GFAP-labelled surface and thickness, in CYCL than in CTRL mice (**Fig. 2D**). The density of microglial cells within the AP and NTS tended to be lower (P=0.054) in CYCL compared to CTRL mice (**Fig. 2D**). Despite these cellular changes observed in histological DVC slices, we did not observe any difference in the expression of genes related to regulation of food intake, plasticity, blood-brain barrier or microglia, astrocytes and inflammation between CTRL and CYCL mice in AP samples isolated by dissection from the brainstem (**Suppl. Table 1**).

### Weight cycling modifies caecal microbiota composition under chow diet

We further explored the impact of weight cycling on gut microbiota by analyzing more precisely the caecal composition of CTRL and CYCL mice under chow diet. Caecal dysbiosis was manifest in CYCL-chow mice when compared to CTRL-chow ones as demonstrated by the principal coordinate analysis using Jaccard distance (**Fig. 3A**). The number of observed species was lower in CYCL than CTRL mice but evenness, showed by the Shannon index, was similar between groups (**Fig. 3B**). At the phylum level, the relative abundance of Bacillota (ex-Firmicutes) was significantly greater while Bacteroidota abundance tended to be lower in the caecum of CYCL than CTRL mice (**Fig. 3C**). Yet, at the genus level, none of the most abundant genera differed between CTRL and CYCL mice (**Fig. 3D**). Only some low abundant genera were altered by weight cycling (13 genera with abundance <0.1%, **Suppl. Table 2**). Finally, we evaluated the SCFA concentration in the caecum, that was significantly greater in CYCL mice compared to CTRL ones, despite the same chow diet, confirming the metabolic alteration of the bacterial community consequently to alternance of WD and chow diets (**Fig. 3E**). Moreover, the relative proportions of the major SCFAs was altered with significant lower proportions of acetate and propionate but a greater proportion of butyrate in CYCL compared to CTRL mice (**Fig. 3E**).

**Figure 3.**
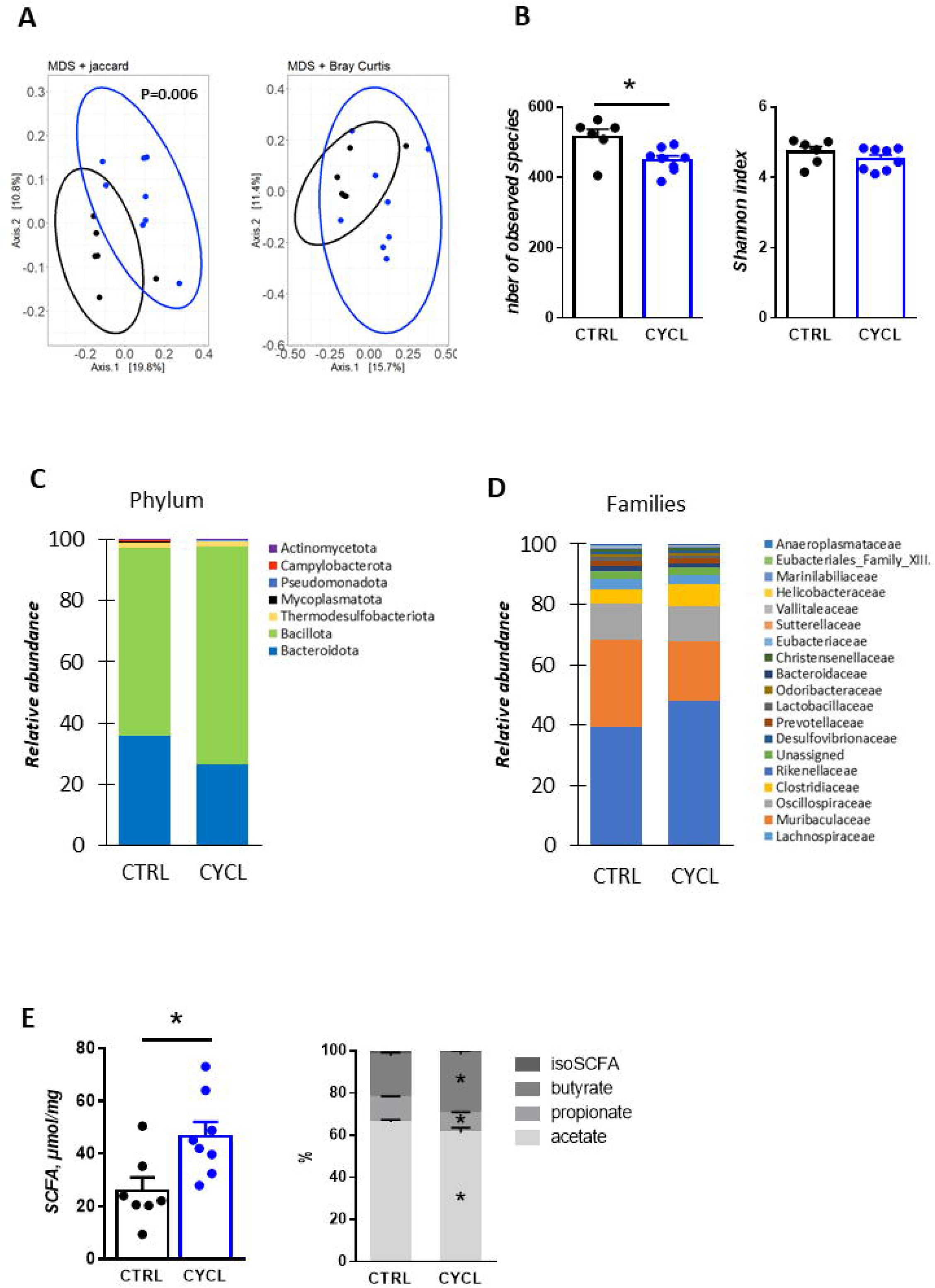
CYCL mice display caecal dysbiosis under chow diet. **(A)** Principal Coordinates Analysis using Jaccard and Bray-Curtis distances of the caecal microbiota of CTRL and CYCL mice under chow diet. **(B)** Number of observed species and Shannon index of the caecal microbiota of CTRL and CYCL mice under chow diet. **(C)** Relative abundances of the different phyla in the caecum of CTRL and CYCL mice under chow diet. **(D)** Relative abundances of the 30 most abundant genera in the caecum of CTRL and CYCL mice under chow diet. **(E)** Caecal total short-chain fatty acids (SCFA) concentration and relative proportions of the different SCFA in the caecum of CTRL and CYCL mice. n=7-8 mice / group, * P<0.05 (Mann-Whitney test). Each dot represents one mouse. Bars are means ± SEM.

### Transferring the microbiota of mice that underwent weight cycling transferred their transient loss of control over WD consumption

The role of the microbiota in regulation of palatable food intake has been recently described (29–31). We thus sought to evaluate if the dysbiosis induced by weight cycling could be at play in the transient loss of control over WD consumption we observed in CYCL mice. Thus, we transferred the caecal microbiota of CYCL mice that had underwent 2 cycles of diet alternance or of age-matched CTRL mice to naïve recipient mice (**Fig. 4A**). We confirmed that the caecal microbiota of donor mice differed, with increased Bacillota and decreased Bacteroidota relative abundances in donor CYCL mice compared to CTRL ones (**Fig. 4B**). Three weeks after transfer, the microbiota of MT-CTRL and MT-CYCL mice tended to differ and the similar differences in phylum relative abundance (increase in Bacillota and decrease in Bacteroidota) observed in the donor mice were observed in the recipient ones (**Fig. 4C**).

**Figure 4:**
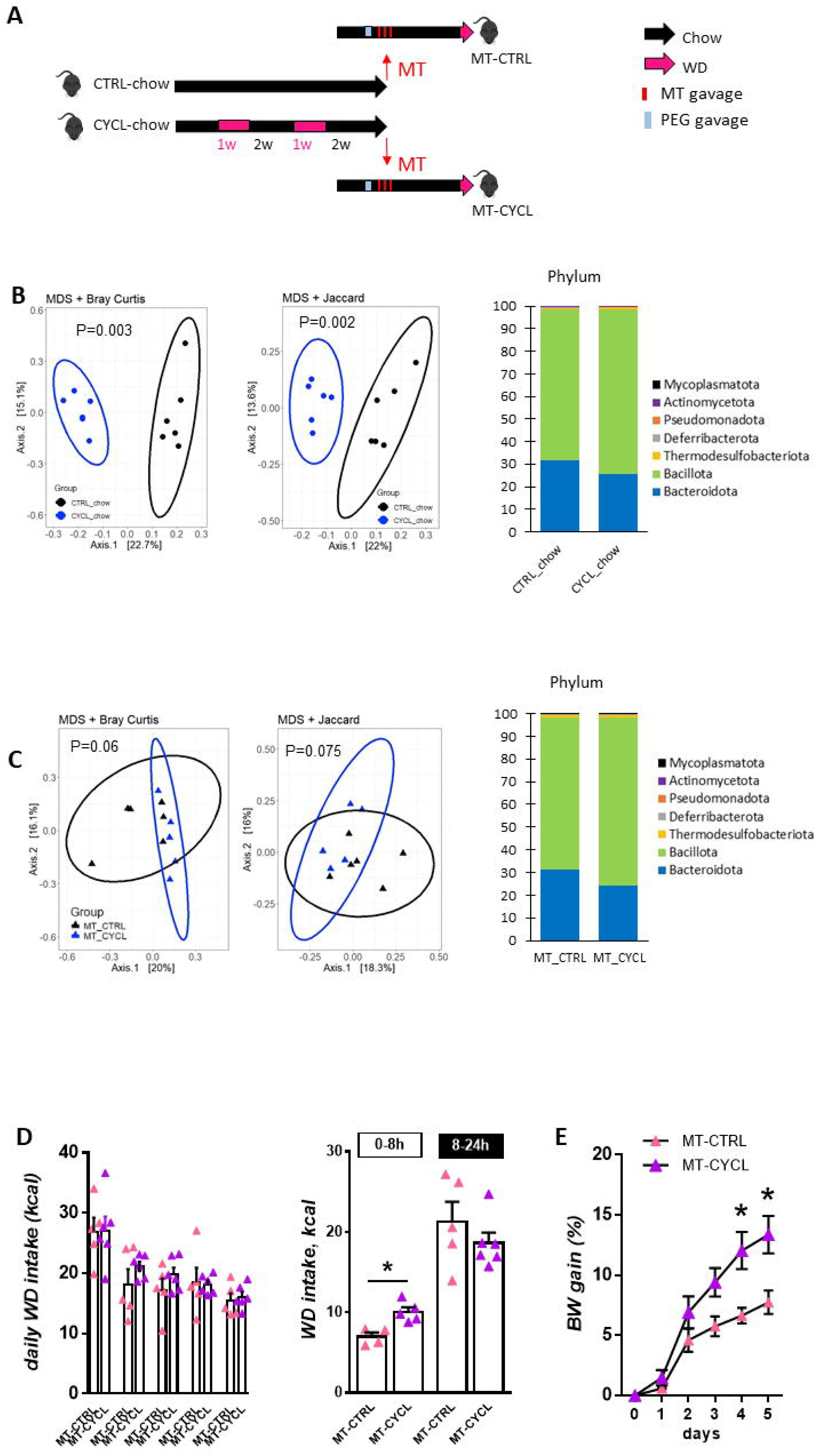
Transferring CYCL-chow microbiota to naïve mice transfers behavioral phenotype. **(A)** Experimental design: Donor mice were submitted to cycles of one week of western diet (WD) followed by 2 weeks of chow diet (CYCL) or left under chow diet (CTRL). They were euthanized under chow and their caecal microbiota transferred on 3 consecutive days (MT transfer) to recipient mice (MT-CYCL and MT-CTRL, respectively) that had been previously prepared by polytethylene glycol (PEG) gavage. Recipient mice were left undisturbed under chow diet for 3 weeks before being either euthanized or switched to WD for 5 days. **(B)** Principal coordinate analysis based on Jaccard and Bray-Curtis distance and relative abundance of phyla of CTRL and CYCL donor mice. **(C)** Principal coordinate analysis based on Jaccard and Bray Curtis distances and relative abundance of phyla in the caecal microbiota of recipient mice (MT-CTRL and MT-CYCL) 3 weeks after microbiota transfer. **(D)** Daily energy intake after WD introduction and energy intake the first 8 and following 16 hours after the switch to WD. **(E)** Body weight gain after WD introduction in recipient mice. * P<0.05 (Mann-Whitney test). n=5-6 mice / group. Each dot represents one mouse. Bars are means ± SEM.

We evaluated the eating behavior response to WD introduction of transferred mice by exposing them to WD for 5 days. As expected, mice exhibited a first 24h-hyperphagia period upon WD introduction, that normalized after 1 day, irrespective of their group (**Fig. 4D**). When we examined more precisely the WD intake during the first 8 hours following WD introduction, MT-CYCL mice ate significantly more WD than MT-CTRL ones (**Fig. 4D**), suggesting a loss of control over WD consumption, as observed in donor mice. Moreover, as observed in donor mice, body weight gain during the 5 days under WD was greater in MT-CYCL mice than MT-CTRL ones (**Fig. 4D**). Noteworthy, eating behavior and body weight gain of transferred mice did not differ under chow diet before WD introduction (**Suppl. Fig. 1**).

## Discussion

This study demonstrates that alternating between high-energy/palatable and standard diets, a model of yoyo effect of recurrent restrictive dieting, remodels the gut microbiota-brain axis towards a profile that is associated with heightened hedonic appetite and risk of weight gain.

To detect potential modifications in eating behavior, we chose to study weight cycling in a mouse model with *ad libitum* access to food. In this model, we show that mice gain and lose weight during WD and chow periods, respectively. In addition, each dietary switch was accompanied by opposite food intake adaptations. A period of transitory hyperphagia was observed during the transition from the chow diet to WD, while a transient hypophagia was measured when switching from WD back to the chow diet. The WD, which is high in sugar and fat, is a highly palatable diet able to stimulate food intake even in the absence of hunger, leading to food overconsumption (32,33). The superior palatability of WD and activation of the mesolimbic system (34) classically explain the increase in food consumption when transitioning to WD. After a few days on WD, food intake normalizes likely due in part to hypothalamic rewiring of POMC neurons (35). When switching from WD to chow diet, transient hypophagia has been previously observed and was associated with changes in hypothalamic and mesolimbic activity (36,37). Indeed, both short-term and prolonged exposure to a palatable diet result in the devaluation of a less palatable diet, reducing the consumption of chow diet. Determining whether these specific changes in food intake patterns when switching diets are the primary drivers of weight cycling in our model remains challenging. In studies that use alternating high-energy/palatable and standard diets to mimic weight cycling in rodents, some authors reported a similar increase in appetite for WD (15), while other did not found this effect (14,38). In these studies, weight cycling was modeled in obese subjects, and weight gain was mainly attributed to reduced locomotor activity and energy expenditure which we did not examined in our study. Therefore, while increased and decreased energy intake might contribute to weight gain and loss in our model, other mechanisms might be involved as well.

Interestingly, WD-induced hyperphagia progressively intensifies during the first few hours of each cycle of WD introduction, followed by a corresponding reduction in WD intake in the subsequent hours. This pattern suggests that components of food reward (i.e. liking, wanting and learning) may be affected in our model. Although this hyperphagic behavior could be attributed to a learned anticipation of WD, as seen as in models of intermittent access to palatable food (39,40), the increase in WD consumption was already evident by the second exposure. Since the reintroduction of WD occurred at a time that could not be predicted by the animals, it rules out the possibility of such learning. The progressive increase in behavioral response to the palatable diet upon re-exposure is also reminiscent of behavioral sensitization observed with repeated drug use (41,42) and may indicate an increased reinforcing value of the WD reward. Interestingly, the drug incentive-sensitization theory is proposed to explain binge-eating disorders, where excessive eating results from excessive food wanting without necessarily increased liking (41,43,44). Furthermore, withdrawal of WD has been shown to elevate stress (45), a state known to exacerbate food wanting and mesolimbic reactivity (46,47). Therefore, repeated WD withdrawal in our model might progressively enhance WD wanting. Finally, despite the escalation in energy intake during the first few hours of renewed WD access, the total energy consumption over the first 24 hours remained stable. This was due to a shift where excessive eating occurred primarily at the beginning of WD introduction, followed by a later reduced WD intake. These data suggest that although food reward valuation and wanting processes might be altered in this model, the homeostatic regulation of food intake is still functional. Therefore, weight maintenance during restrictive yoyo dieting might be impeded not only by metabolic adaptations but also by modified food reward-related processes. This is further emphasized by the increased expression gene involved in the opioidergic and dopaminergic system in the striatum of CYCL mice. The striatum is a component of the mesolimbic system, where these neuromodulatory systems play a crucial role in liking and wanting of food reward (48–51).

In the current study, we found that not only the striatum but also the astroglial barrier between the AP and NTS was affected by alternation between WD and chow diets. The AP and NTS, located in the DVC of the brainstem, integrate signals from the gut and facilitate communication with brain structures involved in eating behavior (52). In this study, we evidenced for the first time a long-lasting effect of WD on astrocyte deployment in the brainstem (53), despite the mice being on a chow diet for the last two weeks. The thicker astroglial barrier in CYCL mice likely serves as a protective mechanism against circulating peripheral signals entering the AP, where the blood brain barrier is leaky (25). Interestingly, we noted a decrease in microglial cells in the AP of CYCL mice fed chow diet. Generally, microglial density increases in response to WD exposure, indicating a pro-inflammatory situation (54). The reduced number of microglial cells may be related to the marked decreased food intake observed in CYCL mice during the second cycle of chow diet. This lower intake likely led to a reduced influx of circulating metabolites, such as fatty acids, which may have tempered the microglial presence and activation within the AP.

In our model, gut microbiota of weight cycling mice was impoverished, a consistent characteristic of WD-chronically fed individuals. Since MT-CYCL mice show heightened hedonic appetite, this could suggest that an impoverished gut microbiota is associated with altered reward processes. Several preclinical studies using microbiota transfer from WD-chronically fed rodents established that these impoverished gut microbiotas alter reward signaling, motivational drive and food intake (55–57). This highlights the suppressive role of gut microbiota on hedonic feeding (30). Our data indicate that yoyo dieting leads to an increased abundance of Bacillota (ex-Firmicutes) at the expense of Bacteroidota. Such change in Bacillota / Bacteroidota ratio is commonly observed in human and mouse models of obesity (58) and increased abundance of Bacillota is associated with increased energy harvesting and fat storage (59,60). In our study, the increase of Bacillota relative abundance was measured in CYCL mice under chow before switching to WD as well as in MT-CYCL. Since Bacillota / Bacteroidota ratio is proposed by some authors to be a potential hallmark for obesity, we could reasonably argue that CYCL-induced dysbiosis in our model could reflect an early risk marker for obesity. However, this finding remains controversial and other authors refute its consideration as a robust marker of obesity-associated dysbiosis (61).

The Bacillota phylum contains many bacteria recognized as SCFA producers (61,62). This is consistent with our data where total SCFA was increased in the caecum of CYCL mice compared to CTRL ones. SCFA have several local effects in the intestine but can also act at distance onto brain functions directly through their passage into the bloodstream or indirectly through modulation of GLP1 and PYY levels, vagus nerve activity or regulation of inflammation (29). SCFA, among others microbiota metabolites, have been shown to regulate reward-related behaviors such as social interactions or drug use (63) as well as eating behaviors (63–65). Our results reveal that the relative proportions of the major SCFAs were altered, with lower proportions of acetate and propionate but a greater proportion of butyrate in CYCL compared to CTRL mice. Since bacteria belonging to the Bacteroidota phylum produce mostly acetate and propionate, and those of the Bacillota phylum have butyrate as their main primary end product (66), these results are consistent with the increased Bacillota / Bacteroidota ratio we observed.

At the genus level, in mice with a history of weight cycling the abundance of several bacteria was significantly decreased (Acetivibrio, Evtepia, Frisingicoccus, Ihubacter, Jutongia, Ruthenibacterium, Tyzzerella, Velocimicrobium) and of others increased (Anaerotaenia, Enteroscipio, Marseillibacter, Muricomes) compared to CTRL mice. While hypotheses could be posited regarding the changes in abundance of these specific bacteria and their consequences on host physiology and eating behavior, these complex interactions are beyond the scope of this study and would require dedicated studies. In addition, the relatively limited overlap in microbial profiles at the genus level between mice and humans (67) is important to acknowledge in the context of translating these results to human health. Further work is thus warranted to understand whether this remodeling at the genus level modifies significantly host physiology and behavior and its relevance to human health.

To conclude, in this study we showed that alternation between high-energy and standard diet durably remodels the gut microbiota towards a profile that is associated with an increase in hedonic appetite and weight gain. Using gut microbiota transfer, we established that this yoyo microbiota signature affects hedonic appetite. Nevertheless, dysbiosis-induced increase in hedonic appetite cannot fully explain weight gain and more work is necessary to fully understand the mechanisms at play in this model; especially regarding the gut microbiota to brain transduction pathways involved in weight cycling and altered eating behavior. However, we believe that this work adds to the comprehension of the interplay between gut microbiota, eating behavior and weight management as collectively, they highlight a putative microbiota signature in individuals prone to weight gain and at risk for hyperphagia of palatable food.

### Study limitations

This study has several limitations that should be considered when interpreting the results. Firstly, we did not implement operant tests to measure reward-related motivational and learning aspects, nor did we assess the hedonic impact of food through taste reactivity tests. Given that changes in the reinforcing value of food are known to predict weight changes in humans, future research should focus on different components of reward, particularly “wanting,” mesolimbic reactivity, and stress within this model to provide a more comprehensive understanding. Additionally, the study was conducted exclusively on male mice, which limits the generalizability of the findings to female mice. The introduction of the WD at the beginning of the light phase may have also influenced the circadian rhythm of the animals, potentially exacerbating weight gain. However, this timing was crucial for identifying changes in hedonic appetite. Future studies should consider these aspects to enhance the robustness and applicability of the findings.

## Acknowledgements

We thank Xufei Zhang for her technical help during animal experiment 2.

## Funding statement

This work was partly funded by INRAE AlimH division and a Groupe Lipides Nutrition grant.

## Conflict of interest

The authors declare no conflict of interest.

## Author contributions

Conceptualization : GB, VD; Methodology : MF, GB, VD; Software : MM; Validation : GB, VD; Formal analysis : MF, GB, VD, MM; Investigation : MF, AS, SR MM, GCP, MM GR, MSH, LLG, CP, ID, VD, GB; Resources : MF, SR, GCP, LLG, ID, VD, GB; Data curation : MF, MM; Writing – original draft : MF, AS, ID, VD, GB; Writing – review & editing : MF, AS, ID, SBF, AB, VD, GB; Visualization : AS, ID, VD, GB; Supervision : VD, GB; Project administration : VD, GB; Funding acquisition : AB, VD, GB

---

**Supplemental Figure 1:**
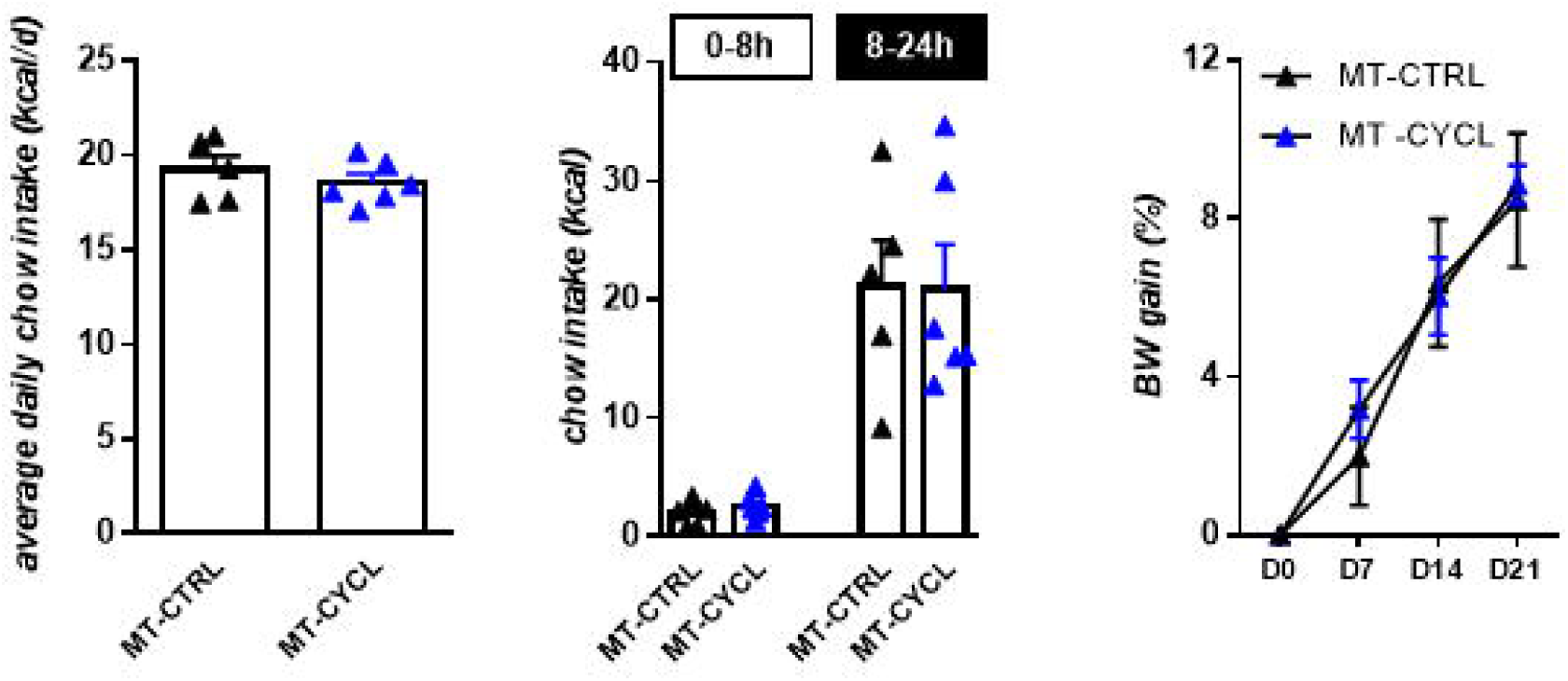
Average daily energy intake, energy intake the last 8 and 16 h before the switch to WD and body weight gain in recipient mice under chow diet.

## Notes

### Competing Interest Statement

The authors have declared no competing interest.

